# Neural signatures of vigilance decrements predict behavioural errors before they occur

**DOI:** 10.1101/2020.06.29.178970

**Authors:** Hamid Karimi-Rouzbahani, Alexandra Woolgar, Anina N. Rich

## Abstract

There are many monitoring environments, such as railway control, in which lapses of attention can have tragic consequences. Problematically, sustained monitoring for rare targets is difficult, with more misses and longer reaction times over time. What changes in the brain underpin these “vigilance decrements”? We designed a multiple-object monitoring (MOM) paradigm to examine how the neural representation of information varied with target frequency and time performing the task. Behavioural performance decreased over time for the rare target (monitoring) condition, but not for a frequent target (active) condition. This was mirrored in the neural results: there was weaker coding of critical information during monitoring versus active conditions. We developed new analyses that can predict behavioural errors from the neural data more than a second before they occurred. This paves the way for pre-empting behavioural errors due to lapses in attention and provides new insight into the neural correlates of vigilance decrements.

## Introduction

When people monitor displays for rare targets, they are slower to respond and more likely to miss those targets relative to frequent target conditions (Wolfe et al., 2005; Warm et al., 2008; Rich et al., 2008). This effect is more pronounced as the time doing the task increases, which is often called a ‘vigilance decrement’. Theoretical accounts of vigilance decrements fall into two main categories. ‘Cognitive depletion’ theories suggest performance drops as cognitive resources are ‘used up’ by the difficulty of sustaining attention under vigilance conditions (Helton et al., 2008; Helton et al., 2011;Warm et al., 2008). In contrast, ‘mind wandering’ theories suggest that the boredom of the task tends to result in insufficient involvement of cognitive resources, which in turn leads to performance decrements (Manly et al., 1999; Smallwood et al., 2006; Young et al., 2002). Either way, there are many real-life situations where such a decrease in performance over time can lead to tragic consequences, such as the Paddington railway disaster (UK, 1999), in which a slow response time to a stop signal resulted in a train moving another 600 meters past the signal into the path of an oncoming train. With the move towards automated and semi-automated systems in many high-risk domains (e.g., power-generation and trains), humans now commonly need to monitor systems for infrequent computer failures or errors. These modern environments challenge our attentional systems and make it urgent to understand the way in which monitoring conditions change the way important information about the task is encoded in the human brain.

To date, most vigilance and rare target studies have used simple displays with static stimuli. Traditional vigilance tasks, inspired by radar operators in WWII (Mackworth, 1948), require participants to respond to infrequent visual events on otherwise blank screens (Temple et al., 2000). Contemporary vigilance tasks, like the Sustained Attention to Response Task (SART), require participants to respond frequently to a rapid stream of static displays and occasionally withhold a response (Rosvold et al., 1956; Rosenberg et al., 2013). However, modern environments (e.g., rail and air traffic control) have additional challenges not encapsulated by these measures. This includes multiple moving objects, potentially appearing at different times, and moving simultaneously in different directions. When an object moves in the space, its neural representation has to be continuously updated so we can perceive the object as having the same identity. Tracking moving objects also requires considerable neural computation: in addition to spatial remapping, for example, we need to predict direction, speed, and the distance of the object to a particular destination. These features cannot be studied using static stimuli; they require objects that shift across space over time. In addition, operators have complex displays requiring selection of some items while ignoring others. We therefore need new approaches to study vigilance decrements in situations that more closely resemble the real-life environments in which humans are now operating. Developing these methods will provide a new perspective on fundamental questions of how the brain implements sustained attention in moving displays, and the way in which monitoring compared with active task involvement changes the encoding of task information. These new methods may also provide avenues to optimise performance in high-risk monitoring environments.

The brain regions involved in maintaining attention over time has been studied using functional Magnetic Resonance Imaging (fMRI), which measures changes in cerebral blood flow (Adler et al., 2001; Benedict et al., 2002; Coull et al., 1996; Gilbert et al., 2006; Johannsen et al., 1997; Ortuno et al., 2002; Perin et al., 2010; Scnell et al., 2007; Sturm et al., 1999; Tana et al., 2010; Thakral et al., 2009; Wingen et al., 2008). These studies compared brain activation in task vs. resting baseline or sensorimotor control (which involved no action) conditions and used univariate analyses to identify regions with higher activation under task conditions. This has the limitation that there are many features that differ between the contrasted (subtracted) conditions, not just the matter of sustained attention. Specifically, this comparison cannot distinguish whether the activation during sustained attention is caused by the differences in the task, stimuli, responses or a combination of these factors. As it is challenging to get sufficient data from monitoring (vigilance) tasks in the scanner, many previous studies used tasks with relatively frequent targets, in which vigilance decrements usually do not occur. However, despite these challenges, Langner et al. (2013) reviewed vigilance neuroimaging studies and identified a network of right-lateralized brain regions including dorsomedial, mid- and ventrolateral prefrontal cortex, anterior insula, parietal and a few subcortical areas that they argue form the core network subserving vigilant attention in humans. The areas identified by Langner et al. (2013) show considerable overlap with a network previously identified as being recruited by many cognitively challenging tasks, the ‘multiple demand’ (MD) regions, which include the right inferior frontal gyrus, anterior insula and intra parietal sulcus (Duncan & Owen, 2000; Duncan, 2010; Fedorenko et al., 2013; Woolgar et al., 2011; Woolgar et al., 2015a; Woolgar et al., 2015b).

Other fMRI studies of vigilance have focused on the default mode network, composed of discrete areas in the lateral and medial parietal, medial prefrontal, and medial and lateral temporal cortices such as posterior cingulate cortex (PCC) and ventral anterior cingulate cortex (vACC), which is thought to be active during ‘resting state’ and less active during tasks (Greicius et al., 2003; Greicius et al., 2009; Raichle et al., 2015). Eichele et al., (2008) suggested that lapses in attention can be predicted by decrease of deactivation of this default mode network. In contrast, Weissman et al. (2006) identified deactivation in the anterior cingulate and right prefrontal regions in pre-stimulus time windows when targets were missed. More recently, Sadaghiani et al. (2015) showed that the functional connectivity between sensory and ‘vigilance-related’ (Cingulo-Opercular) brain areas decreased prior to behavioural misses in an auditory task while the connectivity increased between the same sensory area and the default-mode network. These suggest that modulation of interactions between sensory and vigilance-related brain areas might be responsible for behavioural misses in monitoring tasks.

Detecting changes in brain activation that correlate with lapses of attention can be particularly challenging with fMRI, given that it has poor temporal resolution. Electroencephalography (EEG), which records electrical activity at the scalp, has much better temporal resolution, and has been the other major approach for examining changes in brain activity during sustained attention tasks.

Frequency band analyses have shown that low-frequency alpha (8 to 10.9 Hz) oscillations predict task workload and performance during monitoring of simulated air traffic (static) displays with rare targets, while frontal theta band (4 to 7.9 Hz) activity predicts task workload only in later stages of the experiment (Kamzanova et al., 2014). Other studies find that increases in occipital alpha oscillations can predict upcoming error responses (Mazaheri et al., 2009) and misses (O’Connell et al., 2009) in go/no-go visual tasks with target frequencies of 11% and 9%, respectively. These changes in signal power that correlate with the task workload or behavioural outcome of trials are useful, but provide relatively coarse-level information about what changes in the brain during vigilance decrements.

Understanding the neural basis of decreases in performance over time under vigilance conditions is not just theoretically important, it also has potential real-world applications. In particular, if we could identify a reliable neural signature of attentional lapses, then we could potentially intervene prior to any overt error. For example, with the development of autonomous vehicles, being able to detect when a driver is not engaged, combined with information about a potential threat, could allow emergency braking procedures to be initiated. Previous studies have used physiological measures such as pupil size (Yoss, et al., 1970), body temperature (Molina et al., 2019), skin conductance, blood pressure, etc. (Lohani et al., 2019) to indicate the level of human arousal or alertness, but these lack the fine-grained information necessary to distinguish transient dips from problematic levels of inattention in which task-related information is lost. In particular, we lack detail on how information processing changes in the brain during vigilance decrements. This knowledge is crucial to develop a greater theoretical and practical understanding of how humans sustain vigilance.

In this study, we developed a new task, multiple object monitoring (MOM), which includes key features of real-life situations confronting human operators in high-risk environments. These features include moving objects, varying levels of target frequency, and a requirement to detect and avoid collisions. We recorded neural data using the highly-sensitive method of magnetoencephalography (Baillet, 2017) and used multivariate pattern analyses (MVPA) to detect changes in information encoded in the brain. We used these new approaches to better understand the way in which changes between active and monitoring tasks affects neural processing, including functional connectivity. We then examined the potential for using these neural measures to predict forthcoming behavioural misses based on brain activity.

## Methods

### Participants

We tested twenty-one right-handed participants (10 male, 11 female, mean age = 23.4 years (SD = 4.7 years), all Macquarie University students) with normal or corrected to normal vision. The Human Research Ethics Committee of Macquarie University approved the experimental protocols and the participants gave informed consent before participating in the experiment. We reimbursed each participant AU$40 for their time completing the MEG experiment, which lasted for about 2 hours including setup.

### Apparatus

We recorded neural activity using a whole-head MEG system (KIT, Kanazawa, Japan) with 160 coaxial first-order gradiometers, at a sampling rate of 1000 Hz. We projected the visual stimuli onto a mirror at a distance of 113 cm above participants’ heads while they were in the MEG. An InFocus IN5108 LCD back projection system (InFocus, Portland, Oregon, USA), located outside the magnetically shielded room, presented the dynamically moving stimuli, controlled by a desktop computer (Windows 10; Core i5 CPU; 16 GB RAM; NVIDIA GeForce GTX 1060 6GB Graphics Card) using MATLAB with Psychtoolbox 3.0 extension (Brainard, 1997; Kleiner et al., 2007). We set the refresh rate of the projector at 60 Hz and used parallel port triggers and a photodiode to mark the beginning (dot appearing on the screen) and end (dot disappearing off the screen) of each trial. We recorded participant’s head shape using a pen digitizer (Polhemus Fastrack, Colchester, VT) and placed five marker coils on the head which allowed the location of the head in the MEG helmet to be monitored during the recording-we checked head location at the beginning, half way through and the end of recording. We used a fibre optic response pad (fORP, Current Designs, Philadelphia, PA, USA) to collect responses and an EyeLink 1000 MEG-compatible remote eye-tracking system (SR Research, 1000 Hz monocular sampling rate) to record eye position. We focused the eye-tracker on the right eye of the participant and calibrated the eye-tracker immediately before the start of MEG data recording.

### Task and Stimuli

#### Task summary

The task was to avoid collisions of relevant moving dots with the central object by pressing the space bar if the dot passed a deflection point in a visible predicted trajectory without changing direction to avoid the central object (see Figure 1A; a demo can be found here https://osf.io/c6hy9/). A text cue at the start of each block indicated which colour of dot was relevant for that block. The participant only needed to respond to targets in this colour; dots in the other colour formed distractors. Pressing the button deflected the dot in one of two possible directions (counterbalanced) to avoid collision.

**Figure 1.**
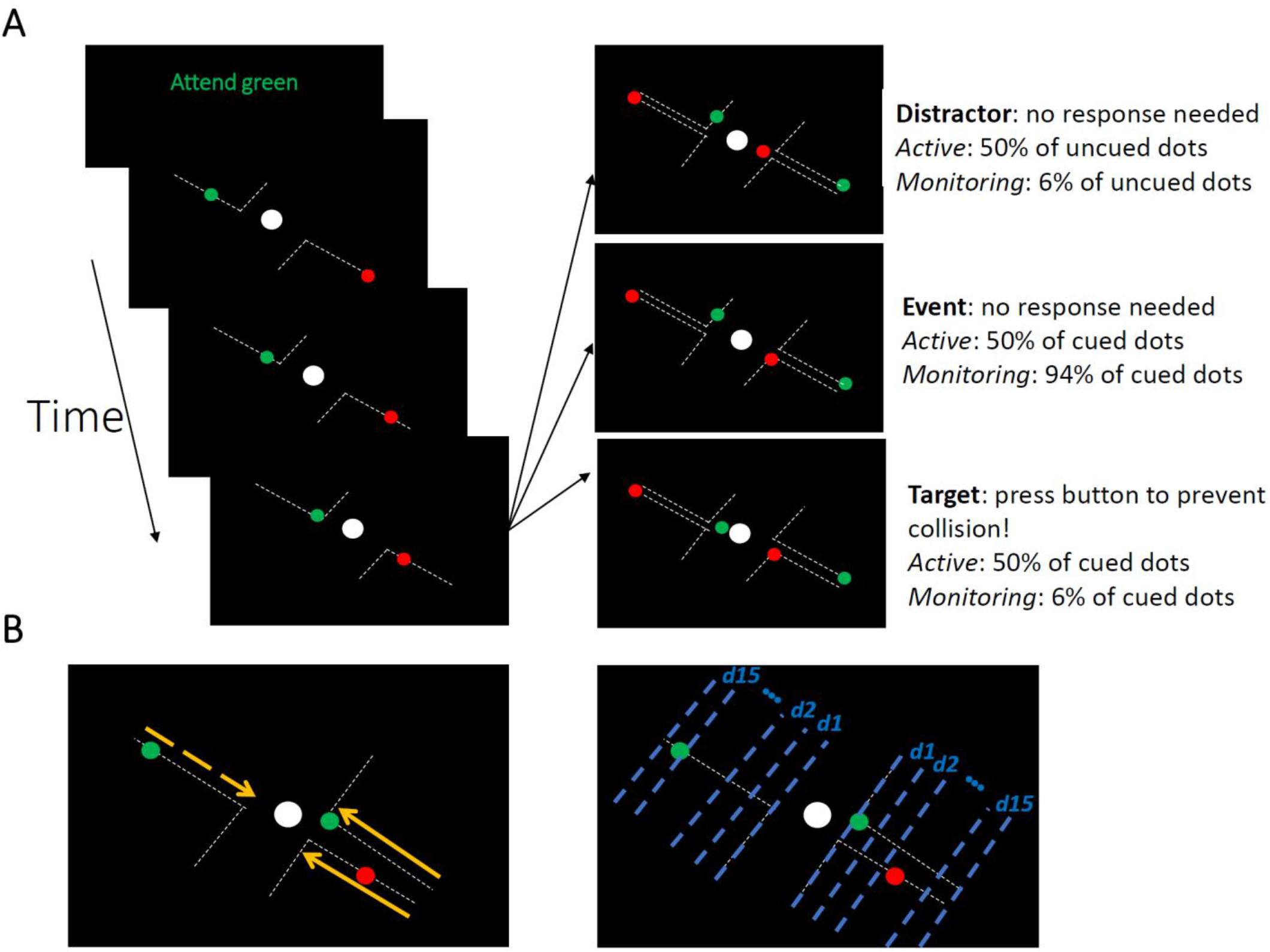
The Multiple Object Monitoring (MOM) task and types of information decoded. (A) At the start of a block, the relevant colour is cued (here, green; distractors in red). Over the on-task period (~30 mins per task condition), multiple dots entered from either direction, each moving along a visible individual trajectory towards the middle object. Only attended dots that failed to deflect along the trajectories at the deflection point required a response (Target: bottom right display). Participants did not need to press the button for the unattended dot (Distractor: top right display) and the dots that kept moving on the trajectories (Event: middle right panel). Each dot took ~1226 ms from appearance to deflection. (B) Direction of approach information (left display: left vs. right as indicated by dashed and solid lines, respectively) and distance information (right display). Note the blue dashed lines and orange arrows were not present in the actual display. A demo of the task can be found here [https://osf.io/c6hy9/].

#### Stimuli

The stimuli were moving dots in one of two colours that followed visible trajectories and covered a visual area of 3.8 × 5 degrees of visual angle (dva; Figure 1A). We presented the stimuli in blocks of 110 s duration, with at least one dot moving on the screen at all times during the 110s block. The trajectories directed the moving dots from two corners of the screen (top left and bottom right) straight towards a centrally presented static “object” (a white dot of 0.25 dva) and then deflected away (either towards the top right or bottom left of the screen; in pathways orthogonal to their *direction of approach*) from the static object at a set distance (the deflection point).

Target dots deviated from the visible trajectory at the deflection point and continued moving towards the central object. The participant had to push the space bar to prevent a ‘collision’. If the response was made before the dot reached the centre of the object, the dot deflected, and this was counted as a ‘hit’. If the response came after this point, the dot continued straight, and this was counted as a ‘miss’, even if they pressed the button before the dot totally passed through central object.

The time from dot onset in the periphery to the point of deflection was 1226±10 (Mean ± SD) milliseconds. Target (and distractor event) dots took 410±10 (Mean ± SD) milliseconds to cross from the deflection point to the collision point. In total, each dot moved across the display for 2005±12 (Mean ± SD) milliseconds before starting to fade away after either deflection or travel through the object. The time delay between the onsets of different dots (ISI) was 1660±890 (Mean ± SD) milliseconds. There were 1920 dots presented in the whole experiment (~56 mins). Each 110 second block contained 64 dots, 32 (50%) in red and 32 (50%) in green, while the central static object and trajectories were presented in white on a black background.

#### Conditions

There were two target frequency conditions. In ‘Monitoring’ blocks, target dots were ~6.2% of cued-colour dots (2 out of 32 dots). In ‘Active’ blocks, target dots were 50% of cued-colour dots (16 out of 32 dots). The same proportion of dots in the non-cued colour failed to deflect; these were distractors (see Figure 1A, top right panel). Participants completed two practice blocks of the Active condition and then completed 30 blocks in the main experiment (15 Active followed by 15 Monitoring or *vice versa*, counterbalanced across participants).

The time between the appearance of target dots varied unpredictably, with distractors and correctly-deflecting dots (events) intervening. In Monitoring blocks, there was an average time between targets of 57.88 (±36.03 SD) seconds. In Active blocks, there was an average time between targets of 7.20 (±6.36 SD) seconds.

#### Feedback

On target trials, if the participant pressed the space bar in time, this ‘hit’ was indicated by a specific tone and deflection of the target dot. There were three types of potential false alarm, all indicated by an error tone and no change in the trajectory of the dot. These were if the participant responded: (1) too early, while the dot was still on the trajectory; (2) when the dot was not a target and had been deflected automatically (‘event’ in Figure 1A, middle right); or (3) when the dot was in the non-cued colour (‘distractor’ in Figure 1A, top right) in any situation. Participants had only one chance to respond per dot; any additional responses resulted in ‘error’ tones. As multiple dots could be on the screen, we always associated the button press to the dot which was closest to the central object.

### Pre-processing

MEG data were filtered online using band-pass filters in the range of 0.03 to 200 Hz and notch-filtered at 50 Hz. We did not perform eye-blink artefact removal because it has been shown that blink artefacts are successfully ignored by multivariate classifiers as long as they are not systematically different between decoded conditions (Grootswagers et al., 2017). We then imported the data into Matlab and epoched them from −100 to 3000 ms relative to the trial onset time. Finally, we down-sampled the data to 200 Hz for the decoding of our two key measures: *direction of approach* and *distance to object* (see below).

### Multivariate pattern analyses (MVPA)

We measured the information contained in the multivariate (multi-sensor) patterns of MEG data by training a linear discriminant analysis (LDA) classifier using a set of training trials from two categories (e.g., for the *direction of approach* measure, this was dots approaching from left vs. right, see below). We then tested to see whether the classifier could predict the category of an independent (left-out) set of testing data from the same participant. We used a 10-fold cross-validation approach, splitting the data into training and testing subsets. Specifically, we trained the LDA classifier on 90% of the trials and tested it on the left-out 10% of the trials. This procedure was repeated 10 times each time leaving out a different 10% subset of the data for testing (i.e., 10-fold cross validation).

We decoded two major task features from the neural data: (1) the *direction of approach* (left vs. right); and (2) the distance of each moving dot from the centrally fixed object (*distance to object*), which correspond to visual (retinal) information changing over time. Our interest was in the effect of selective attention (attended vs. unattended) and Target Frequency conditions (Active vs. Monitoring) on the neural representation of this information, and how the representation of information changed on trials when participants missed the target.

We decoded left vs. right *directions of approach* (as indicated by yellow arrows in Figure 1B) every 5 ms starting from 100 ms before the appearance of the dot on the screen to 3000 ms later. Please note that as each moving dot is considered a trial, trial time windows (epochs) overlapped for 62.2% of trials. In Monitoring blocks, 1.2% of target trials overlapped (two targets were on the screen simultaneously but lagged relative to one another). In Active blocks, 17.1% of target trials overlapped.

For the decoding of *distance to object*, we split the trials into the time windows corresponding to 15 equally spaced distances of the moving dot relative to the central object (as indicated by blue lines in Figure 1B), with distance 1 being closest to the object, and 15 being furthest away (the dot having just appeared on the screen). Next, we collapsed (concatenated) the MEG signals from identical distances (splits) across both sides of the screen (left and right), so that every distance included data from dots approaching from both left and right side of the screen. This concatenation ensures that distance information decoding is not affected by the *direction of approach*. Finally, we trained and tested a classifier to distinguish between the MEG signals (a vector comprising data from all MEG sensors, concatenated over all time points in the relevant time window), pertaining to each pair of distances (e.g., 1 vs. 2) using a leave-one-out cross-validation procedure. We obtained classification accuracy for all possible pairs of distances (105 combinations of 15 distances). To obtain a single decoding value per distance, we averaged the 14 classification values that corresponded to that distance against other 14 distances. For example, the final decoding accuracy for distance 15 was an average of 15 vs. 14, 15 vs. 13, 15 vs. 12 and so on until 15 vs. 1. We repeated this procedure for our main Target Frequency conditions (Active vs. Monitoring), Attention conditions (attended vs. unattended) and Time on Task (first and last five blocks of each task condition, which are called early and late blocks here, respectively). This was done separately for *correct* and *miss* trials and for each participant separately.

### Informational connectivity analysis

To evaluate possible modulations of brain connectivity between the attentional networks of the frontal brain and the occipital visual areas, we used a simplified version of our recently developed RSA-based connectivity analysis (Goddard et al., 2016; Karimi-Rouzbahani, 2018; Karimi-Rouzbahani et al., 2019). Specifically, we evaluated the informational connectivity, which measures the similarity of distance information between areas, across our main Target Frequency conditions (Active vs. Monitoring), Attention conditions (attended vs. unattended) and Time on Task (first and last five blocks of each task condition, which are called early and late blocks here, respectively). This was separately done for *correct* and *miss* trials, using representational dissimilarity matrices (RDM; Kriegeskorte et al., 2008). To construct the RDMs, we decoded all possible combinations of distances from each other yielding a 15 by 15 cross-condition classification matrix, for each condition separately. We obtained these matrices from peri-occipital and peri-frontal areas to see how the manipulation of Attention, Target Frequency and Time on Task modulated the correlation of information (RDMs) between those areas on *correct* and *miss* trials. We quantified connectivity using Spearman’s rank correlation of the matrices obtained from those areas, only including the lower triangle of the RDMs (105 decoding values). To avoid bias when comparing the connectivity on *correct* vs. *miss* trials, the number of trials were equalized by subsampling the *correct* trials to the number of *miss* trials and repeating the subsampling 100 times before finally averaging them for comparison with *miss* trials.

### Error data analysis

Next, we asked what information was coded in the brain when participants missed targets. To study information coding in the brain on *miss* trials, where the participants failed to press the button when targets failed to automatically deflect, we used our recently-developed method of error data analysis (Woolgar et al., 2019). Essentially, this analysis asks whether the brain represents the information similarly on *correct* and *miss* trials. For that purpose, we trained a classifier using the neural data from a proportion of *correct* trials (i.e., when the target dot was detected and manually deflected punctually) and tested on both the left-out portion of the *correct* trials (i.e., cross-validation) and on the *miss* trials. If decoding accuracy is equal between the *correct* and *miss* trials, we can conclude that information coding is maintained on *miss* trials as it is on *correct* trials. However, if decoding accuracy is lower on *miss* trials than on *correct* trials, we can infer that information coding differs on *miss* trials, consistent with the change in behaviour. Since *correct* and *miss* trials were visually different after the deflection point, we only used data from before the deflection point.

For these error data analyses, the number of folds for cross-validation were determined based on the proportion of *miss* to *correct* trials (number of folds = number of miss trials/number of correct trials). This allowed us to test the trained classifiers with equal numbers of *miss* and *correct* trials to avoid bias in the comparison.

### Predicting behavioural performance from neural data

We developed a new method to predict, based on the most task-relevant information in the neural signal, whether or not a participant would press the button for a target dot in time to deflect it on a particular trial. This method includes three steps, with the third step being slightly different for the left-out testing participant vs. the other 20 participants. First, for every participant, we trained 105 classifiers using ~80% of correct trials to discriminate the 15 distances. Second, we tested those classifiers using half of the left-out portion (~10%) of the correct trials, which we called validation trials, by simultaneously accumulating (i.e., including in averaging) the accuracies of the classifiers at each distance and further distances as the validation dot approached the central object. The validation set allowed us to determine a decision threshold for predicting the outcome of each testing trial: whether it was a *correct* or *miss* trial. Third, we performed a second-level classification on testing trials which were the other half (~10%) of the left-out portion of the correct trials and the miss trials, using each dot’s accumulated accuracy calculated as in the previous step. Accordingly, if the testing dot’s accumulated accuracy was ***higher*** than the decision threshold, it was predicted as *correct*, otherwise *miss*. For all participants, except for the left-out testing one, the decision threshold was chosen from a range of ***multiples*** (0.1 to 4 in steps of 0.1) of the standard deviation below the accumulated accuracy obtained for the validation set on the second step. For determining the optimal threshold for the testing participant, however, instead of a range of multiples, we used the average of the best performing multiples (i.e., the one which predicted the behavioural outcome of the trial more accurately) obtained from the other 20 participants. This avoided circularity in the analysis.

To give more detail on the second and third steps, when the validation/testing dots were at distance #15, we averaged the accuracies of the 14 classifiers trained to classify dots at distance #15 from all other distances. Accordingly, when the dot reached distance #14, we also included and averaged accuracies from classifiers which were trained to classify distance #14 from all other distances leading to 27 classifier accuracies. Therefore, by the time the dot reached distance #1, we had 105 classifier accuracies to average and predict the behavioural outcome of the trial. Every classifier’s accuracies were either 1 or 0 corresponding to correct or incorrect classification of dot’s distance, respectively. Note that accumulation of classifiers’ accuracies, as compared to using classifier accuracy on every distance independently, provides a more robust and smoother classification measure for deciding on the label of the trials. The validation set, which was different from the testing set, allowed us to set the decision threshold based on the validation data within each subject and from the 20 participants and finally test our prediction classifiers on a separate testing set from the 21^st^ individual participant, iteratively. The optimal threshold was 1.54 (± 0.2) times the SD below the decoding accuracy on the validation set across participants.

### Eye-tracking data analysis

To see if we could use a less complicated physiological measure to obtain information about the processing of visual information, and to check that the decoding we observed was not just due to eye movements, we repeated the above decoding analyses using the eye-tracking data. Specifically, instead of the MEG sensor data, we decoded the information about the *direction of approach* and *distance to object* using x-y coordinates of the right eye fixation provided by the eye-tracker. All other aspects of the analysis were identical to the ‘error data analysis’ section. If we observe a similar decoding of information using the eye-tracking data, it would mean that we could use eye-tracking, which is a less expensive and more feasible approach for prediction of errors, instead of MEG. If the prediction from the MEG decoding was stronger than that of the eye tracking, it would mean that there was information in the neural signal over and above any artefact associated with eye movement.

### Statistical analyses

To determine the evidence for the null and the alternative hypotheses, we used Bayes analyses as implemented by Krekelberg (https://klabhub.github.io/bayesFactor/) based on Rouder et al. (2012).

We used standard rules for interpreting levels of evidence (Lee and Wagenmakers, 2014; Dienes, 2014): Bayes factors of >10 and <1/10 were interpreted as strong evidence for the alternative and null hypotheses, respectively, and >3 and <1/3 were interpreted as moderate evidence for the alternative and null hypotheses, respectively. We interpreted the Bayes factors which fell between 3 and 1/3 as reflecting insufficient evidence either way.

Specifically, for the behavioural data, we asked whether there was a difference between Active and Monitoring conditions in terms of miss rates and reaction times. Accordingly, we calculated the Bayes factor as the probability of the data under alternative (i.e., difference) relative to the null (i.e., no difference) hypothesis in each block separately. In the decoding, we repeated the same procedure to evaluate the evidence for the alternative hypothesis of a difference between decoding accuracies across conditions (e.g. Active vs. Monitoring and Attended vs. Unattended) vs. the null hypothesis of no difference between them, at every time point/distance. To evaluate evidence for the alternative of above-chance decoding accuracy vs. the null hypothesis of no difference from chance, we calculated the Bayes factor between the distribution of actual accuracies obtained and a set of 1000 random accuracies obtained by randomising the class labels across the same pair of conditions (null distribution) at every time point/distance.

To evaluate the evidence for the alternative of main effects of different factors (Attention, Target Frequency and Time on Task) in decoding, we used Bayes factor ANOVA (Rouder et al., 2012). This analysis evaluates the evidence for the null and alternative hypothesis as the ratio of the Bayes factor for the full model ANOVA (i.e., including all three factors of Target Frequency, Attention and the Time on Task) relative to the restricted model (i.e., including the two other factors while excluding the factor being evaluated). For example, for evaluating the main effect of Time on Task, the restricted model included Attention and Target Frequency factors but excluded the factor of Time on Task.

The priors for all Bayes factor analyses were determined based on Jeffrey-Zellner-Siow priors (Jeffreys, 1961; Zellner and Siow, 1980) which are from the Cauchy distribution based on the effect size that is initially calculated in the algorithm using a *t*-test (Rouder et al., 2012). The priors are data-driven and have been shown to be invariant with respect to linear transformations of measurement units (Rouder et al., 2012), which reduces the chance of being biased towards the null or alternative hypotheses.

## Results

### Behavioural data: The MOM task evokes a reliable vigilance decrement

In the first 110 second experimental block of trials (i.e., excluding the two practice blocks), participants missed 29% of targets in the Active condition and 40% of targets in the Monitoring condition. However, the number of targets in any single block is necessarily very low for monitoring conditions (for a single block, there are 16 targets for Active but only 2 targets for Monitoring). The pattern does become more robust over blocks, and Figure 2A shows the miss rates changed over time in different directions for the Active vs. Monitoring conditions. For Active blocks, miss rates decreased over the first five blocks and then plateaued at ~17%. For Monitoring, however, miss rates increased throughout the experiment: by the final block, these miss rates were up to 76% (but again, the low number of targets in Monitoring mean that we should use caution in interpreting the results of any single block alone). There was evidence that miss rates were higher in the Monitoring than Active conditions from the 4th block onwards (BF > 3; Figure 2A). Participants’ reaction times (RTs) on correct trials also showed evidence of vigilance decrements, increasing over time under Monitoring but decreasing under Active task conditions (Figure 2B). There was evidence that reaction times were slower for Monitoring compared with Active from the sixth block onwards (BF > 3, except for Block #11). The characteristic pattern of increasing miss rates and slower RTs over time in the Monitoring relative to the Active condition validates the MOM task as effectively evoking vigilance decrements.

**Figure 2.**
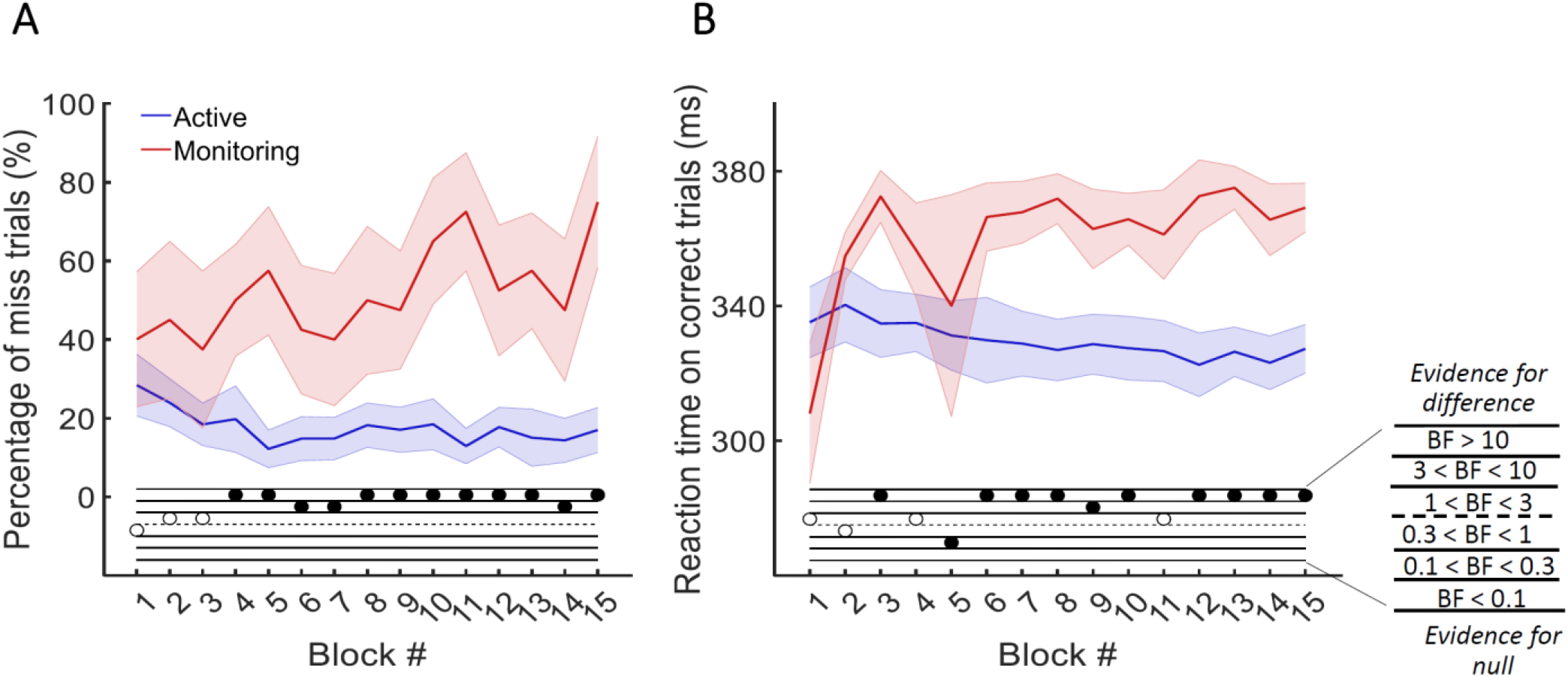
Behavioural performance on the MOM task. The percentage of *miss* trials (A), and *correct* reaction times (B), as a function of block. Thick lines show the average across participants (shading 95% confidence intervals) for Active (blue) and Monitoring (red) conditions. Each block lasted for 110 seconds and had either 16 (Active) or 2 (Monitoring) targets out of 32 cued-colour and 32 non-cued colour dots. Bayes Factors are shown in the bottom section of each graph: Filled circles show moderate/strong evidence for either hypothesis and empty circles indicate insufficient evidence when evaluating the contrast between Active and Monitoring conditions.

### Neural data: Decoding different aspects of task-related information

With so much going on in the display at one time, we first needed to verify that we can successfully decode the major aspects of the moving stimuli, relative to chance. The full data figures and details are presented in Supplementary Materials: We were able to decode both *direction of approach* and *distance to object* relative to chance from MEG signals (see Supplementary Figure 1). Thus, we can turn to our main question about how these representations were affected by the Target Frequency, Attention and Time on Task.

### The neural correlates of the vigilance decrement

As the behavioural results showed (Figure 2), the difference between Active and Monitoring conditions increased over time, showing the greatest difference during the final blocks of the experiment. To explore the neural correlates of these vigilance decrements, we evaluated information processing in the brain during the first five and last five blocks of each task (called early and late blocks, respectively) and the interactions between the Target Frequency, Attention and the Time on Task using a 3-way Bayes factor ANOVA as explained in *Methods*.

Effects of Target Frequency on *direction of approach* information *Direction of approach* information is a very clear visual signal (‘from the left’ vs ‘from the right’) and therefore is unlikely to be strongly modulated by other factors, except perhaps whether the dot was in the cued colour (Attended) or the distractor colour (could be ignored: Unattended). There was strong evidence for a main effect of Attention (Figure 3A; BF > 10, Bayes factor ANOVA, cyan dots) starting from 265ms and lasting until dots faded. This is consistent with maintenance of information about the attended dots and attenuation of the information about unattended dots (Supplementary Figure 1A). The large difference in coding attributable to attention remained for as long as the dots were visible.

**Figure 3.**
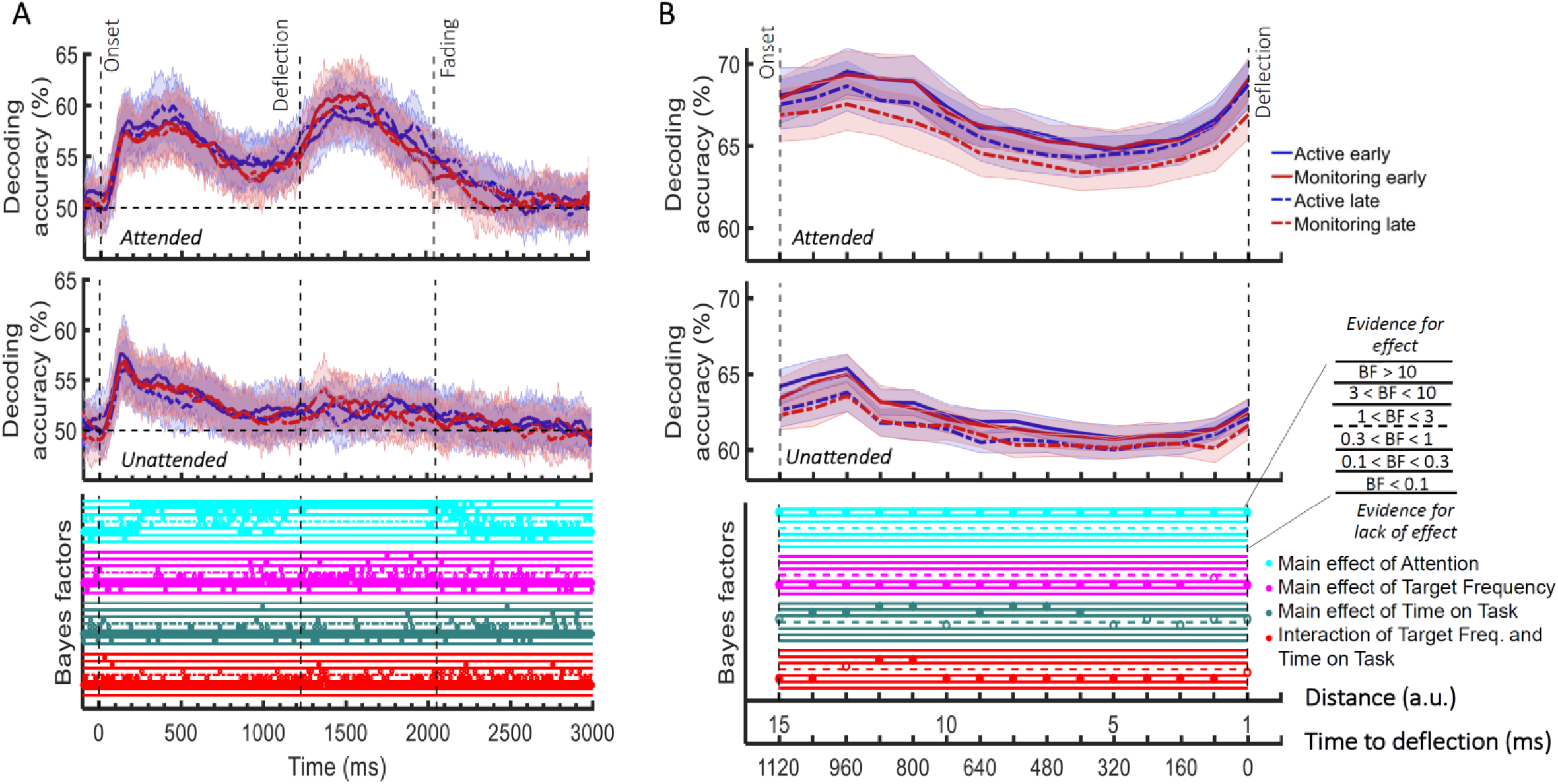
Impact of different conditions and their interactions on information processing on *correct* trials (all trials except those in which a target was missed or there was a false alarm). (A) Decoding of *direction of approach* information (less task-relevant). The horizontal dashed line refers to theoretical chance-level decoding (50%). Upper graph: Attended dot; Lower graph: Unattended (‘distractor’) dot. (B) Decoding of *distance to object* information (most task-relevant) and their Bayesian evidence for main effects and interactions. Thick lines show the average across participants (shading 95% confidence intervals). Vertical dashed lines indicate critical times in the trial. Bayes Factors are shown in the bottom section of each graph: Filled circles show moderate/strong evidence for either hypothesis and empty circles indicate insufficient evidence. Main effects and interactions of conditions calculated using Bayes factor ANOVA analysis. Cyan, pink, green and red dots indicate the main effects of Attention, Target frequency, Time on Task and the interaction between Target frequency and Time on Task, respectively. The results of Bayes factor analysis (i.e. the main effects of the three conditions and their interactions) are from the same 3-way ANOVA analysis and therefore identical for attended and unattended panels. Early = data from the first 5 blocks (~10 minutes). Late = data from the last 5 blocks (~10 minutes).

In contrast, there was no sustained main effect of Target Frequency on the same *direction of approach* coding (0.1 < BF < 0.3; Bayes factor ANOVA, Figure 3A, pink dots). For the majority of the epoch there was moderate evidence for the null hypothesis (BF < 1/3). The sporadic time points with a main effect of Target Frequency, observed a few times before the deflection (3 < BF < 10), likely reflect noise in the data as there is no clustering. Recall that we only focus on timepoints prior to deflection, as after this point there are visual differences between Active and Monitoring, with more dots deflecting in the Monitoring condition.

There was also no sustained main effect of the Time on Task on information about the *direction of approach* (0.1 < BF < 0.3; Bayes factor ANOVA, green dots; Figure 3A). There were no sustained 2-way or 3-way interactions between Attention, Target Frequency and Time on Task (BF < 1; Bayes factor ANOVA). Note that the number of trials used in the training and testing of the classifiers were equalized across the 8 conditions and equalled the minimum available number of trials across those conditions shown in Figure 3. Therefore, the observed effects cannot be attributed to a difference in the number of trials across conditions.

### Effects of Target Frequency on critical *distance to object* information

The same analysis for the representation of the task-relevant *distance to object* information showed strong evidence for a main effect of Attention (BF > 10; Bayes factor ANOVA) at all 15 distances, moderate or strong evidence for a main effect of Time on Task (BF > 3; Bayes factor ANOVA) at eight of the earlier distances, and an interaction between Time on Task and Target Frequency at two of these distances (Figure 3B). There was more decoding for attended than unattended dots (compare top and bottom panels of Figure 3B). The main effect of Time on Task reflected decreased decoding in later blocks (compare dashed lines to solid lines in Figure 3B). Finally, the interaction between Target Frequency and Time on Task can be seen when comparing the solid to the dashed lines in blue and red colours, separately, and suggests a bigger decline in decoding in Monitoring compared to Active conditions. Note that as there was moderate evidence for no interaction between Attention and Target Frequency or between Attention and Time on Task (0.1 < BF < 0.3, 2-way Bayes factor ANOVA) or simultaneously between the three factors (BF < 0.1, 3-way Bayes factor ANOVA), we do not show those statistical results in the figure.

Together, these results suggest that while vigilance conditions had little or no impact on coding of the *direction of approach*, they did impact the critically task-relevant information about the distance of the dot from the object. Coding of this information declined as the time on the task increased and this effect was more pronounced when the target events happened infrequently.

### Is brain connectivity modulated by Attention, Target Frequency and the Time on Task?

Using graph-theory-based univariate connectivity analysis, it has been recently shown that the connectivity between relevant sensory areas and “vigilance-related” cognitive areas changes prior to lapses in attention (behavioural misses; Sadaghiani et al., 2015). Therefore, we asked whether vigilance decrements across the time course of our task corresponded to changes in multi-variate connectivity, which tracks information transfer, between frontal attentional networks and sensory visual areas. Specifically, we asked whether there were changes in *information exchange* between these conditions. We used a simplified version of our method of RSA-based informational connectivity to evaluate the (Spearman’s rank) correlation between distance information RDMs across the peri-frontal and peri-occipital electrodes (see *Methods*; Goddard et al., 2016; Figure 4A).

**Figure 4.**
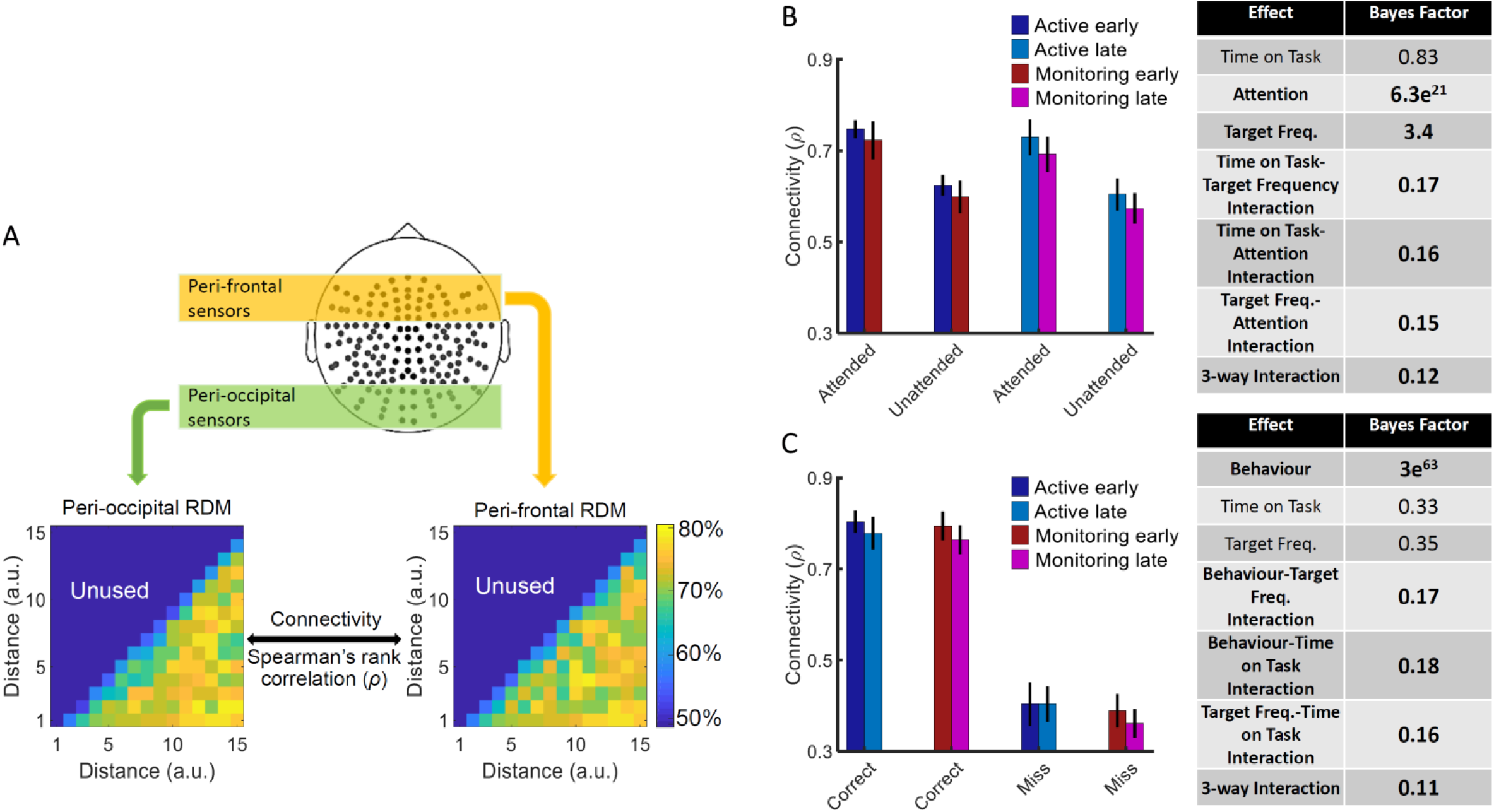
Relationship between informational connectivity and Attention, Target Frequency, Time on Task and the behavioural outcome of the trial (i.e., correct vs. miss). (A) Calculation of connectivity using Spearman’s rank correlation between RDMs obtained from the peri-frontal and peri-occipital sensors as indicated by colored boxes, respectively. RDMs include decoding accuracies obtained from testing the 105 classifiers trained to discriminate different *distance to object* categories. (B) Connectivity values for the eight different conditions of the task and the results of three-way Bayes factor ANOVA with factors Time on Task (early, late), Attention (attended, unattended) and Target Frequency (active, monitoring), using only correct trials. (C) Connectivity values for the Active and Monitoring, Early and Late blocks of each task for *correct* and *miss* trials (attended condition only) and the result of Bayes factor ANOVA with factors Target Frequency (Active, Monitoring), Time on Task (early, late) and behavioural outcome (correct, miss) as inputs. Number of trials are equalized across conditions in B and C separately. Bars show the average across participants (error bars 95% confidence intervals). Bold fonts indicate moderate or strong evidence for either the effect or the null hypothesis.

Results showed strong evidence (Bayes factor ANOVA, BF = 6.3e^21^) for higher informational connectivity for Attended compared to Unattended trials, and moderate evidence for higher connectivity in Active compared to Monitoring conditions (Bayes factor ANOVA, BF = 3.4; Figure 4B). There was insufficient evidence to determine whether there was a main effect of Time on Task (Bayes factor ANOVA, BF = 0.83). There was moderate evidence for no 2-way and 3-way interactions between the three factors (Bayes factor ANOVA, 2-way Time on Task-Target Frequency: BF = 0.17; Time on Task-Attention: BF = 0.16; Target Frequency-Attention: BF = 0.15; their 3-way interactions BF = 0.12). These results suggest that Monitoring conditions and trials in which the dots are in the distractor (unattended) colour, in which the attentional load is low, result in less informational connectivity between occipital and frontal brain areas compared to Active conditions and attended trials, respectively. This is consistent with a previous study (Alnaes et al., 2015), which suggested that large-scale functional brain connectivity depends on the attentional load, and might underpin or accompany the decrease in information decoding across the brain in these conditions (Figure 3B).

We also compared the connectivity for the correct vs. miss trials (Figure 4C). This analysis was performed only for attended condition as there are no miss trials for unattended condition, by definition. There was strong evidence for less (almost half) connectivity on *miss* compared to *correct* trials (Bayes factor ANOVA, BF = 3e^63^). There was insufficient evidence to determine the effects of the Time on Task or Target Frequency (Bayes factor ANOVA, BF = 0.33 and BF = 0.35, respectively) and moderate evidence for a lack of 2-way and 3-way interactions between the three factors (Bayes factor ANOVA, Behaviour-Target Frequency: BF = 0.17; Behaviour-Time on Task: BF = 0.18; Target Frequency-Time on Task: BF = 0.16; their 3-way interactions BF = 0.11). Weaker connectivity between occipital and frontal areas could have led to the behavioural misses observed in this study (Figure 1) as was previously reported in an auditory monitoring task using univariate graph-theoretic connectivity analyses (Sadaghiani et al., 2015), although, of course, these are correlational data and so we cannot make any strong causal inferences. These results cannot be explained by the number of trials as they are equalized across the 8 conditions in each of the analyses separately.

### Can we use the neural data to predict behavioural errors before they occur?

#### Is neural information processing different on *miss* trials?

The results presented in Figure 3, which used only *correct* trials, showed changes due to target frequency to the representation of task-relevant information when the task was performed successfully. We next move on to our second question, which is whether these neural representations change when overt behaviour *is* affected, and therefore, whether we can use the neural activity as measured by MEG to predict behavioural errors before they occur. We used our method of error data analysis (Woolgar et al., 2019) to examine whether the patterns of information coding on miss trials differed from correct trials (see *Methods*). For these analyses we used only attended dots, as unattended dots do not have behavioural responses, and we matched the total number of trials in our implementation of correct and miss classification.

First, we evaluated the processing of the less relevant information - the *direction of approach* measure (Figure 5A). The results for *correct* trials provided information dynamics very similar to the attended condition in Figure 3A, except for higher overall decoding, which is explained by the inclusion of the data from the whole experiment (15 blocks) rather than just the five early and late blocks (note the number of trials is still matched to miss trials).

**Figure 5.**
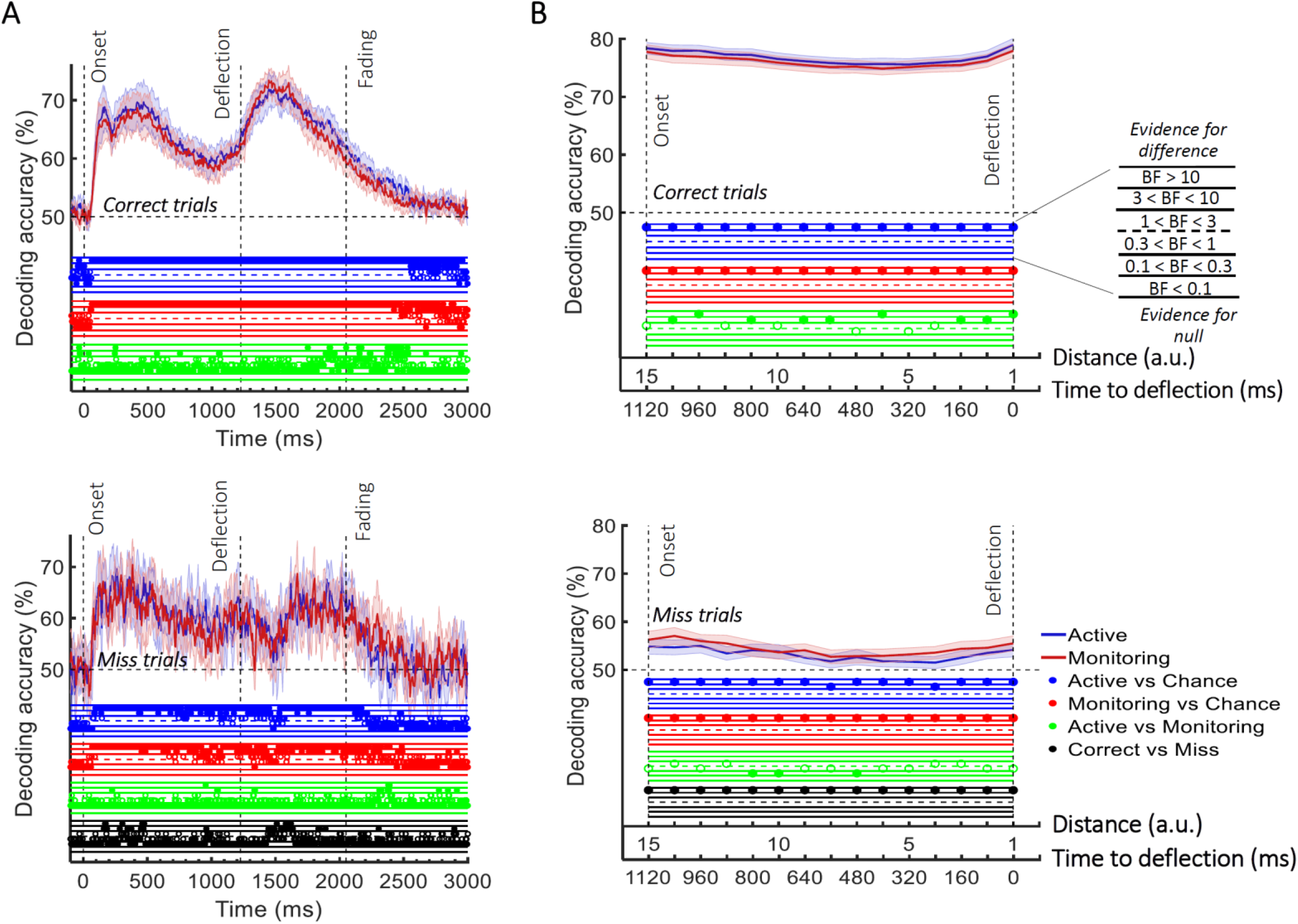
Decoding of information on *correct* vs. *miss* trials. (A) Decoding of *direction of approach* information (less task-relevant). (B) Decoding of *distance to object* information (most task-relevant). The horizontal dashed lines refer to chance-level decoding. Top panels: Decoding using correct trials; Bottom panels: Decoding using miss trials. In both top and bottom panels, the classifiers were trained on *correct* trials and tested on (left out) *correct* and all *miss* trials, respectively. Thick lines show the average across participants (shading 95% confidence intervals). Vertical dashed lines indicate critical events in the trial. Bayes Factors are shown in the bottom section of each graph: Filled circles show moderate/strong evidence for either hypothesis and empty circles indicate insufficient evidence. They show the results of Bayes factor analysis when evaluating the difference of the decoding values from chance for Active (blue) and Monitoring (red) conditions separately, the comparison of the two conditions (green) and the comparison of correct and miss trials (black). Note that for the comparison of correct and miss trials, Active and Monitoring conditions were averaged separately.

Active and Monitoring conditions did not show any time windows of sustained difference (BF < 0.3). However, when the classifiers were tested on *miss* trials, from onset to deflection, the pattern of information dynamics were different, even though we had matched the number of trials.

Specifically, while the level of information was comparable to *correct* trials with spurious instances (but no sustained time windows) of difference (BF > 3 as indicated by black dots) before 500 ms, decoding traces were much noisier for *miss* trials with more variation across trials and between nearby time points (Figure 5A). Note that after the deflection, the visual signal is different for correct and miss trials, so the difference between their decoding curves (BF > 3) is not meaningful. These results suggest a noisier processing of *direction of approach* information for the missed dots compared to correctly deflected dots.

We then repeated the same procedure on the processing of the most task-relevant *distance to object* information on *correct* vs. *miss* trials (Figure 5B). Although on *correct* trials, the distance information for both Active and Monitoring conditions was well above chance (77%; BF > 10), for *miss* trials, the corresponding distance information was only just above chance (55%; BF > 10 for all distances except one). The direct comparison revealed that distance information dropped considerably on *miss* trials compared to *correct* trials (Figure 5; Black dots; BF > 10 across all distances; Active and Monitoring results were averaged for correct and miss trials separately before Bayes analyses). This is consistent with less representation of the crucial information about the distance from the object preceding a behavioural miss.

#### Can we predict behavioural errors using neuroimaging?

Finally, we asked whether we could use this information to predict the behavioural outcome of each trial. To do so, we developed a new method that classified trials based on their behavioural outcomes (correct vs. miss) by asking how well a set of classifiers, pre-trained on correct trials, would classify the distance of the dot from the target (see *Methods*; Figure 6A). To achieve this, we used a second-level classifier which labelled a trial as correct or miss based on the average accumulated accuracies obtained for that dot at every distance from the first-level decoding classifiers which were trained on *correct* trials (Figure 6A and 6B; see *Methods*). If the accumulated accuracy for the given dot at the given distance was less than the average accuracy obtained from testing on the validation set minus a specific threshold (based on standard deviation), the testing dot (trial) was labelled as *correct*, otherwise *miss*. As Figure 6B shows, there was strong evidence (BF > 10) that decoding accuracy of distances was higher for *correct* than *miss* trials with the inclusion of more classifier accuracies as the dot approached from the corner of the screen towards the centre with a multiple of around 1.5 as threshold (Figure 6C). This clear separation of accumulated accuracies for *correct* vs. *miss* trials allowed us to predict with above-chance accuracy the behavioural outcome of the ongoing trial (Figure 6D). To find the optimal threshold for each participant, we evaluated the thresholds used for all other participants except for a single testing participant for whom we used the average of the best thresholds that led to highest prediction accuracy for other participants. This was ~1.5 standard deviation below the average accuracy on the other participants’ validation (correct trial) sets (Figure 6C).

**Figure 6.**
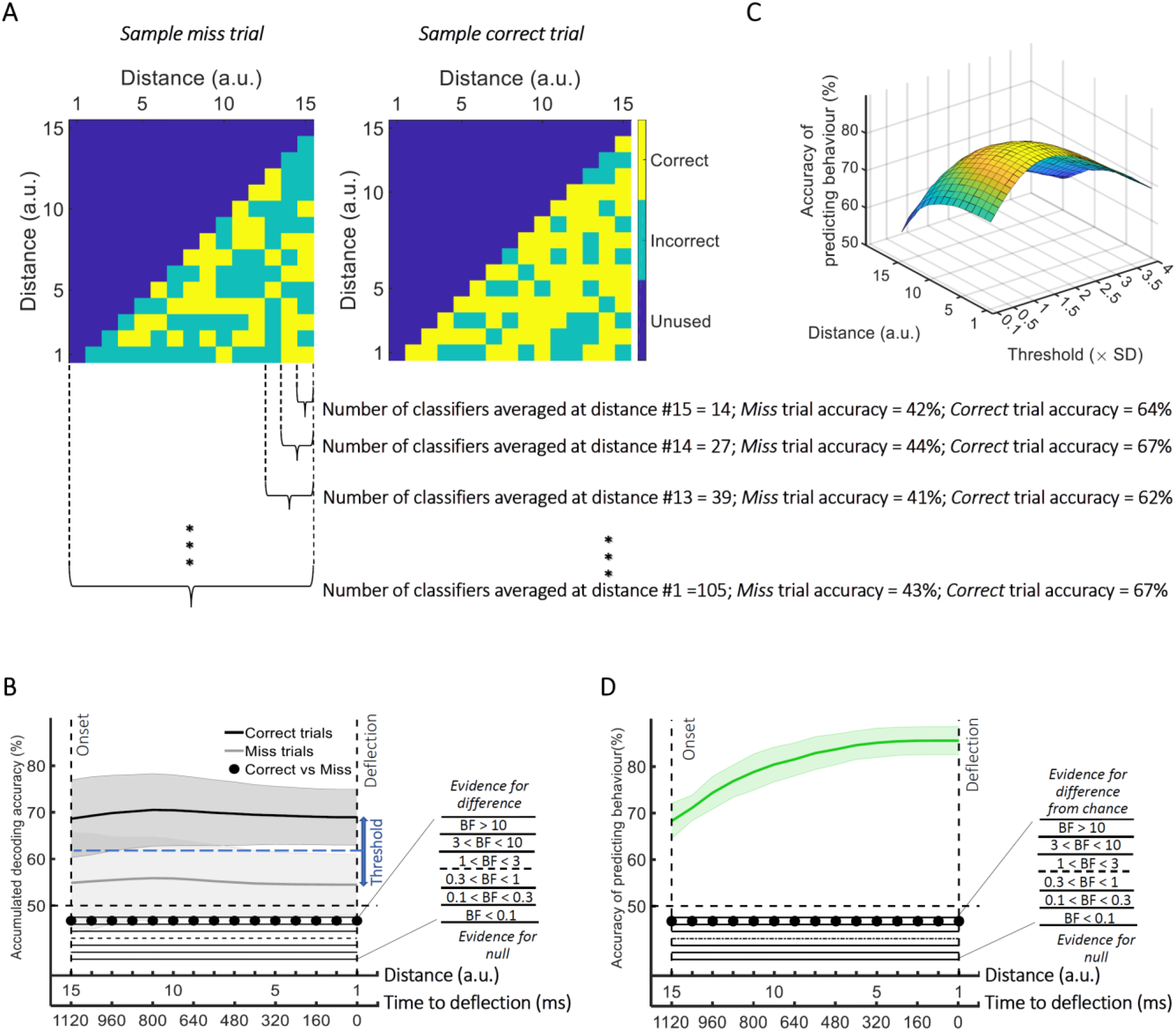
Prediction of behavioural outcome (*correct* vs. *miss*) trial-by-trial using decoding of *distance to object* information. (A) Sample classifiers’ accuracies (correct or incorrect classification of current distance as indicated by colors) for a *miss* (left panel; average accuracy ~= 43% when the dot reached the deflection point) and a *correct* trial (right panel; average accuracy ~=67% at the deflection point). The classifiers were trained on the data from *correct* trials and tested on the data from *correct* and *miss* trials. For the *miss* trials, around half the classifiers classified the dot’s distance incorrectly by the time it reached the deflection point. (B) Accumulation of classifiers’ accuracies over decreasing dot distances/time to deflection. This shows stronger information coding of the crucial *distance to object* information on the *correct* trials over *miss* trials. A variable threshold used in (C) is shown as a blue dashed line. (C) Prediction of behavioural outcome as a function of threshold and distance using a second-level behavioural outcome classification. Results show highest prediction accuracies on the participant set at around the threshold of 1.5 (see *Methods*), increasing at closer distances. (D) Accuracy of predicting behavioural outcome for the left-out participant using the threshold obtained from all the other participants as function of distance/time from the deflection point. Results showed successful (~=70%) prediction of behavioural outcome of the trial as early as 80 ms after stimulus appearance. Thick lines and shading refer to average and one standard deviation around the mean across participants, respectively. Bayes Factors are shown in the bottom section of each graph: Filled circles show moderate/strong evidence for either hypothesis and empty circles indicate insufficient evidence (black dots under B and D).

The prediction accuracy of behavioural outcome was above chance level (68% vs. 50%; BF > 10) even when the dot had only been on the screen for 80ms, which corresponds to our furthest distance #15 (1200ms prior to deflection point; Figure 6D). The accuracy increased to 85% as the dot approached the centre of the screen, with ~80% accuracy with still 800 ms to go before required response. Importantly, the prediction algorithm showed generalisable results across participants; the threshold for decision obtained from the other participants could predict the accuracy of an independent participant’s behaviour using only their neural data.

Please note that the results presented so far were from *correct* and *miss* trials and we excluded early, late and wrong-colour *false alarms* to be more specific about the error type. However, the false alarm results (collapsed across all three types of false alarms) were very similar (Supplementary Figure 2) to those of the missed trials (Figure 5): noisy information about the *direction of approach* and at-chance information about the *distance to object*. This may suggest that both miss and false alarm trials are caused by a similar impaired processing of information, or at least captured similarly by our decoding methods. The average number of miss trials was 58.17 (±21.63 SD) and false alarm trials was 65.94 (±21.13 SD; out of 1920 trials).

### Can we decode direction and distance information from eye-tracking data?

To see whether we could decode information about the dot motion using only the eye-tracking data, we repeated the same error data analysis as above, but this time using the 2-dimensional signals (i.e., corresponding to the x-y coordinates of the gaze location) provided by the eye-tracker (Görgen et al., 2018). The decoding of *direction of approach* from *correct* trials showed above-chance information (Supplementary Figure 3A) starting from 455 and 460 ms post-stimulus onset for the Active and Monitoring conditions (BF > 10), respectively. The information on *miss* trials was noisier but showed a similar pattern. The correct and miss trials only showed moderate evidence (3 < BF < 10) for difference in the span from 310 ms to 490 ms. This suggests that participants moved their eyes differently for the dots approaching from opposite directions, which is not unexpected (and observed in the eye-tracking fixation points data). Although the dynamics of this decoding over time is different to the neural decoding, in line with visually evoked information decoding studies (VanRullen, 2007; Karimi-Rouzbahani et al., 2017), the eye-movement data do hold enough information to decode the *direction of approach*.

In contrast, for the crucial *distance to object* measure, although the eye-tracking data showed above-chance values at a few distances (BF > 10; Supplementary Figure 3B), most were very close to chance and much lower than those obtained from the neural data (cf. Figure 5B; BF > 10 for the difference between decoding of neural vs eyetracking data for correct trials; indicated by black dots in Supplementary Figure 3A). Only for the decoding for *miss* vs. *correct* trials was there any evidence (moderate) for similarity between neural and eyetracking data (0.1 < BF < 0.3; black dots; Supplementary Figure 3B). Note that *distance to object* data collapses across identical distances from the left and right sides of the screen, which avoids the potential confound of eye-movements data driving the classifier for this crucial distance measure.

## Discussion

This study developed new methods to gain insights into how attention, the frequency of target events, and the time doing a task affect the representation of task information in the brain. Our new multiple object monitoring (MOM) task evoked reliable vigilance decrements in both accuracy and reaction time in a situation that more closely resembles real-life modern tasks than classic vigilance tasks. By using the sensitive analysis method of MVPA, we were able to test information coding across task conditions to evaluate the neural correlates of vigilance decrements. We also developed a novel informational brain connectivity method, which allowed evaluation of the correlation between information coding across peri-occipital and peri-frontal areas in different task conditions, to investigate the brain connectivity under different levels of attention, target frequency and the time on the task. Finally, we utilised our recent error data analysis to predict forthcoming behavioural misses with high accuracy. In the following sections, we explain each of the four contributions in detail and compare them with relevant literature.

First, the MOM task includes key features of real-world monitoring situations that are not usually part of other vigilance tasks (e.g., Mackworth, 1948; Temple, 2000; Rosvold et al., 1956; Rosenberg et al., 2013), and the results show clear evidence of vigilance decrements. Behavioural performance, measure with both reaction time and accuracy, deteriorated over time in monitoring (infrequent target) relative to active (frequent target) conditions. These vigilance decrements demonstrate that the MOM task can be used to explore vigilance in situations more closely resembling modern environments, namely involving moving stimuli and selection of relevant from irrelevant information, giving a useful tool for future research.

Second, the high sensitivity of MVPA to extract information from neural signals allowed us to investigate the temporal variations in processing as the experiment progressed. The manipulation of attention showed a strong overall effect with enhanced representation of both the less important *direction of approach* and the most task-relevant *distance to object* information for cued dots, regardless of how frequent the targets were (Figure 3). The improved representation of information under attention extends previous findings from us and others (Woolgar et al., 2015b; Goddard et al., 2019; Nastase et al., 2017) to moving displays, in which the participants monitor multiple objects simultaneously.

The manipulation of target frequency showed that when participants only had to respond infrequently, modelling real-life monitoring situations, the neural coding of crucial information about the task dropped, correlating with the decrease in behavioural performance (i.e., vigilance effects in both accuracy and RT; Figure 2). This suggests that when people monitor for rare targets, they process or encode the relevant information less effectively as the time passes relative to conditions in which they are actively engaged in completing the task. Several previous studies have examined the neural correlates of vigilance decrements using univariate analyses (for a review see Langner et al. (2013)). However, univariate analyses fail to capture widespread but subtle differences of patterns between conditions across distant brain networks. One recent study utilized the sensitivity of MVPA to extract task-relevant and task-irrelevant information under sustained attention (Megan et al., 2015). In this case, however, the aspects of information were similar in identity (i.e. high-level visual categories of face and scenes) and switched their attentional role (i.e. attended vs. unattended) across the experiment, which makes it difficult to see whether (if at all) vigilance decrements would differentially affect encoding of different aspects of information depending on their relevance to the task. To address this issue, here we not only switched the task-relevance of information across the experiment to replicate the attentional effect of that study (i.e. cued/un-cued dots), but we also studied two aspects of the dot motion information that varied in importance for carrying out the task (i.e., *direction of approach* and *distance to object*) with unchanging roles across the experiment. While switching between dot colours showed the effect of attention, with greater representation of the cued dots over uncued dots, the relevance of the *direction of approach* and the *distance to object* did not vary. The less relevant direction information was unaffected by target frequency, whereas the coding of the critical task-relevant distance information correlated with the decrease in behavioural performance over time. This is relevant to theories of vigilance, by demonstrating that the task-relevance of information might be a major factor in whether vigilance decrements occur.

It is important to note that previous studies have tried other physiological/behavioural measures to determine participants’ vigilance or alertness, such as pupil size (Yoss et al., 1970), response time variability (Rosenberg et al., 2013), blood pressure and thermal energy (Lohani et al., 2019) or even body temperature (Molina et al., 2019). We used highly-sensitive analysis of neuroimaging data so that we could address two questions that could not be answered using these more general vigilance measures. Our approach allowed us to test for changes in the way information is processed in the brain, particularly testing for differences in the impact of monitoring on the relevance of the information, rather than whether the participants were vigilant and alert in general. Moreover, we could also investigate how relevant and less relevant information was affected by the target frequency and time on the task, which could explain the behavioural vigilance decrement observed in many previous studies (e.g., Dehais et al., 2019; Wolfe et al., 2005; Wolfe et al., 2007; Kamzanova et al., 2014; Ishibashi et al., 2012). We tested our methods also on the eye-tracking data and found that the critical task-relevant information change under monitoring conditions could not be replicated based on eye-movements, demonstrating the benefit of the neural approach.

Third, our information-based brain connectivity method showed weaker connectivity between the peri-frontal attentional network and the peri-occipital visual areas of the brain in the unattended and monitoring conditions (Figure 4), where participants encountered fewer targets relative to the other conditions. We also observed less connectivity between the same areas on *miss* vs. *correct* trials, which might explain the behavioural outcome of the trials. Most previous neuroimaging studies have used univariate brain connectivity analyses, which are prone to missing existing functional connectivity across areas when encountering low-amplitude activity on individual sensors (Anzellotti & Coutanche, 2018; Basti et al., 2020). The method we used here evaluated the correlation between representational dissimilarity matrices, which has provided high-dimensional information about *distance to object*, obtained from multiple sensors across the brain areas. This makes the analysis more sensitive to capturing subtle connectivity and also aligns with a major recent shift in literature from univariate to multivariate informational connectivity analyses (Goddard et al., 2016; Goddard et al., 2019; Karimi-Rouzbahani et al., 2019; Karimi-Rouzbahani, 2017; Anzellotti & Coutanche, 2018; Basti et al., 2020).

Fourth, building upon our recently-developed method of error analysis (Woolgar et al., 2019), we were able to predict forthcoming behavioural misses before the response was given. This method only used *correct* trials for training, which makes its implementation plausible for real-world situations since we usually have plenty of correct trials and only few miss trials (i.e., cases when the railway controller diverts the trains correctly vs. misses and a collision happens). In our study, the method showed a large decline in the crucial task-relevant (i.e., *distance to object*) information decoding on *miss* vs. *correct* trials but less decline in the less task-relevant information (i.e., *direction of approach*). A complementary analysis allowed the prediction of behaviourally missed trials as soon as the stimulus appeared on the screen (after ~80 ms), which was ~1200 ms before the time of response. This method was generalisable across participants, with the decision threshold for trial classification other participants’ data successful in predicting errors for a left-out participant. A number of previous studies have shown that behavioural performance could be correlated with aspects of brain activity even before the stimulus onset (Eichele et al., 2008; Weissman et al., 2006; Sadaghiani et al., 2015). This can be crucial for many high-risk environments, including semi-autonomous car driving and railway control. Those studies have explained the behavioural errors by implicit measures such as less deactivation of the default-mode network, reduced stimulus-evoked sensory activity (Weissman et al., 2006; Eichele et al., 2008) and even the connectivity between sensory and vigilance-related/default-mode brain areas (Sadaghiani et al., 2015). It would be informative, however, if they could show how (if at all) the processing of task-relevant information is disrupted in the brain and how this might lead to behavioural errors. To serve an applied purpose, it would be ideal if there was a procedure to use those neural signatures to predict behavioural outcomes. Only two previous studies have approached this goal. Sadaghiani et al. (2015) and Dehais et al. (2019) reported maximum prediction accuracies of 63% and 72% (with adjusted chance levels of 55% and 59%, respectively), far lower than what we have obtained here (up to 85% with a chance level of 50%), suggesting our method accesses more relevant neural signatures of vigilance decrements, or is more sensitive in discriminating these. The successful prediction of an error from neural data more than a second in advance of the impending response provides a promising avenue for detecting lapses of attention before any consequences occur.

Current explanations for vigilance effects generally fall into two categories: mind-wandering and cognitive overload. In the first, the low cognitive demands of monitoring tasks result in mind-wandering and then, when a response is required, there are insufficient resources dedicated to the task (e.g., malleable attention theory (Manly et al., 1999; Smallwood et al., 2006; Young et al., 2002)). In the second, the demands of sustaining attention depletes cognitive resources over time leading to insufficient resources and increased errors in later stages of the task (e.g., Helton et al., 2008; Helton et al., 2011; Warm et al., 2008). There are several previous observations of decreased functional connectivity during mind wandering (Chou et al., 2017; Kucyi et al., 2018; van Son et al., 2019), which our informational connectivity results broadly replicate. For example, Chou et al. (2017) reported a decrease in functional connectivity between visual and sensorimotor and in turn to frontal brain areas in later stages of a resting-state mind-wandering fMRI study in which participants were instructed to draw their mind to specific but broad sets of thoughts. In another study, using EEG-fMRI, von Son et al. (2019) found reduced functional connectivity between the dorsolateral PFC, dorsal anterior cingulate cortex (ACC), and posterior parietal regions, namely the “executive control network”, when participants counted and reported their number of inhales and episodes of mind wandering. Our finding of a decrease in higher order cognitive (peri-frontal) and sensory (peri-occipital) areas in later (compared with early) stages of the experiment is broadly consistent with these findings, but we are unable to distinguish whether this is due to mind wandering or the depletion of cognitive resources, as in our task either is a plausible explanation for the effect.

The overall goal of this study was to understand how neural information processing of dynamic displays were affected by attention and target frequency, and whether reliable changes in behaviour over time could be predicted on the basis of neural patterns. We observed that the neural representation of critically relevant information in the brain decreases over time, especially when targets are infrequent. This neural representation was particularly poor on trials where participants missed the target. We used this observation to predict behavioural outcome of individual trials, and showed that we could accurately predict behavioural outcome more than a second before action was needed. These results provide new insights about how vigilance decrements impact information coding in the brain and propose an avenue for predicting behavioural errors using novel neuroimaging analysis techniques.

## Acknowledgments

This work was funded by an Australian Research Council (ARC) Discovery Project grant to A.N.R and A.W. (DP170101780). A.W. was supported by an ARC Future Fellowship (FT170100105) and MRC intramural funding SUAG/052/G101400. We thank Denise Moerel, Mark Wiggins, Jeremy Wolfe and William Helton for contributions to an earlier design of the MOM task.

## Supplementary Material

### Supplementary figure 1 shows the same decoding results as presented in Figure 3 but evaluated against chance-level decoding (50%)

Our first analysis was to verify that our analyses could decode the important aspects of the display, relative to chance, given the overlapping moving stimuli. Here, we give the detailed results of this analysis.

We started with the information about the *direction of approach* (top left or bottom right of screen) which is a strong visual signal but not critical to the task decision. From 95 ms post-stimulus onset onwards, this visual information could be decoded from the MEG signal for all combinations of the factors: Attended and Unattended dots, both Target Frequency conditions (Active, Monitoring), and both our Time on Task durations (Early - first 5 blocks; Late - last 5 blocks; all BF > 10, different from chance).

All conditions were decodable above chance until at least 385 ms post-stimulus onset (BF > 3; Supplementary Figure 1A), which was when the dots came closer to the centre, losing their visual difference. There was a rapid increase in information about the *direction of approach* between 50 ms to 150 ms post-stimulus onset, consistent with an initial forward sweep of visual information processing (VanRullen, 2007; Karimi-Rouzbahani et al., 2017; Karimi-Rouzbahani et al., 2019). For attended dots only (but regardless of the Target Frequency or Time on Task), the information then increased again before the deflection time, and remained different from chance until 1915 ms post-stimulus onset, which is just before the dot faded (Supplementary Figure 1A). The second rise of decoding, which was more pronounced for the attended dots, could reflect the increasing relevance to the task as the dot approached the crucial deflection point, but it could also be due to higher visual acuity in foveal compared to peripheral areas of the visual field. The decoding peak observed after the deflection point for the attended dots, was most probably caused by the large visual difference between the deflection trajectories for the dots approaching from the left vs. right side of the screen (see the deflection trajectories in Figure 1A).

The most task-relevant feature of the motion is the distance between the moving dot and the central object, with the deflection point of the trajectories being the key decision point. We therefore tested for decoding of distance information (*distance to object,* see *Methods*). There was a brief increase in decoding of *distance to object* for attended dots across the other factors (Target Frequency and Time on Task) between the 15^th^ and 10^th^ distances and for the unattended dots across the other factors between 15^th^ and the 12^th^ distances. This corresponds to the first 400 ms for the attended dots and the first 240 ms for the unattended dots after the onset (Supplementary Figure 1B). Distance decoding then dropped somewhat before ascending again as the dot approached the deflection point. The second rise of decoding, which was more pronounced for the attended dots, could reflect the increasing relevance to the task as the dot approached the crucial deflection point, but it could also be due to higher visual acuity in foveal compared to peripheral areas of the visual field. There was strong evidence that decoding of distance information for all conditions was greater than chance (50%, BF > 10) across all 15 distance levels (Supplementary Figure 1B).

**Supplementary Figure 1.**
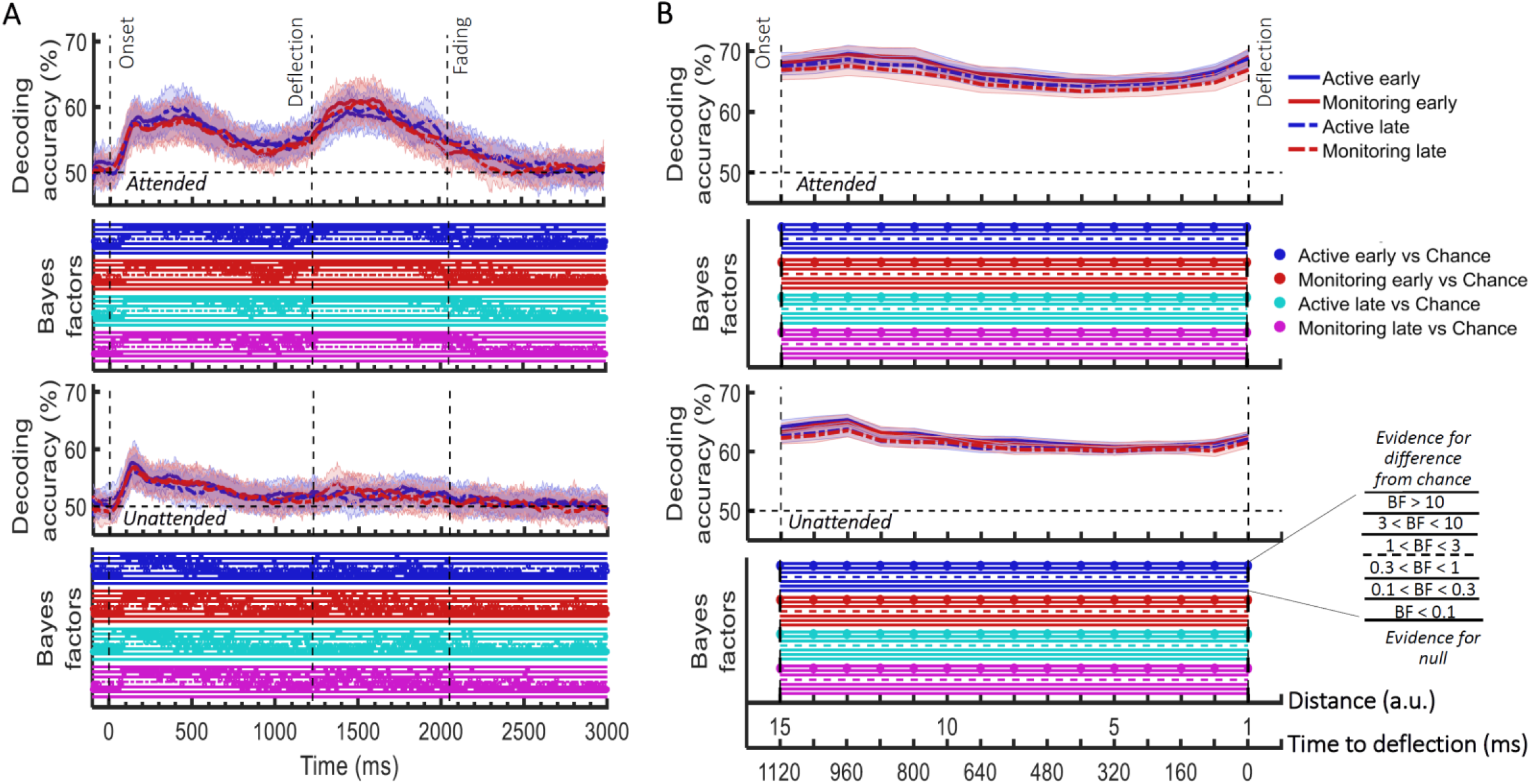
Impact of different conditions in the *direction of approach* (A) and *distance to object* (B) information coding and their Bayesian evidence for difference from chance. (A) Decoding of *direction of approach* information (less task-relevant). The horizontal dashed line refers to chance-level decoding. Upper graph: Attended colour dot; Lower graph: Unattended (‘distractor’) colour dot. (B) Decoding of *distance to object* information (most task-relevant). Thick lines show the average across participants (shading 95% confidence intervals). Vertical dashed lines indicate critical times in the trial. Bayes Factors are shown in the bottom section of each graph: Filled circles show moderate/strong evidence for either hypothesis and empty circles indicate insufficient evidence. They show the results of Bayes factor analysis when evaluating the difference of the decoding values from chance as explained in *Methods*. Early = data from the first 5 blocks (~10 minutes). Late = data from the last 5 blocks (~10 minutes).

### Supplementary figure 2 shows the data for false alarm trials

**Supplementary Figure 2.**
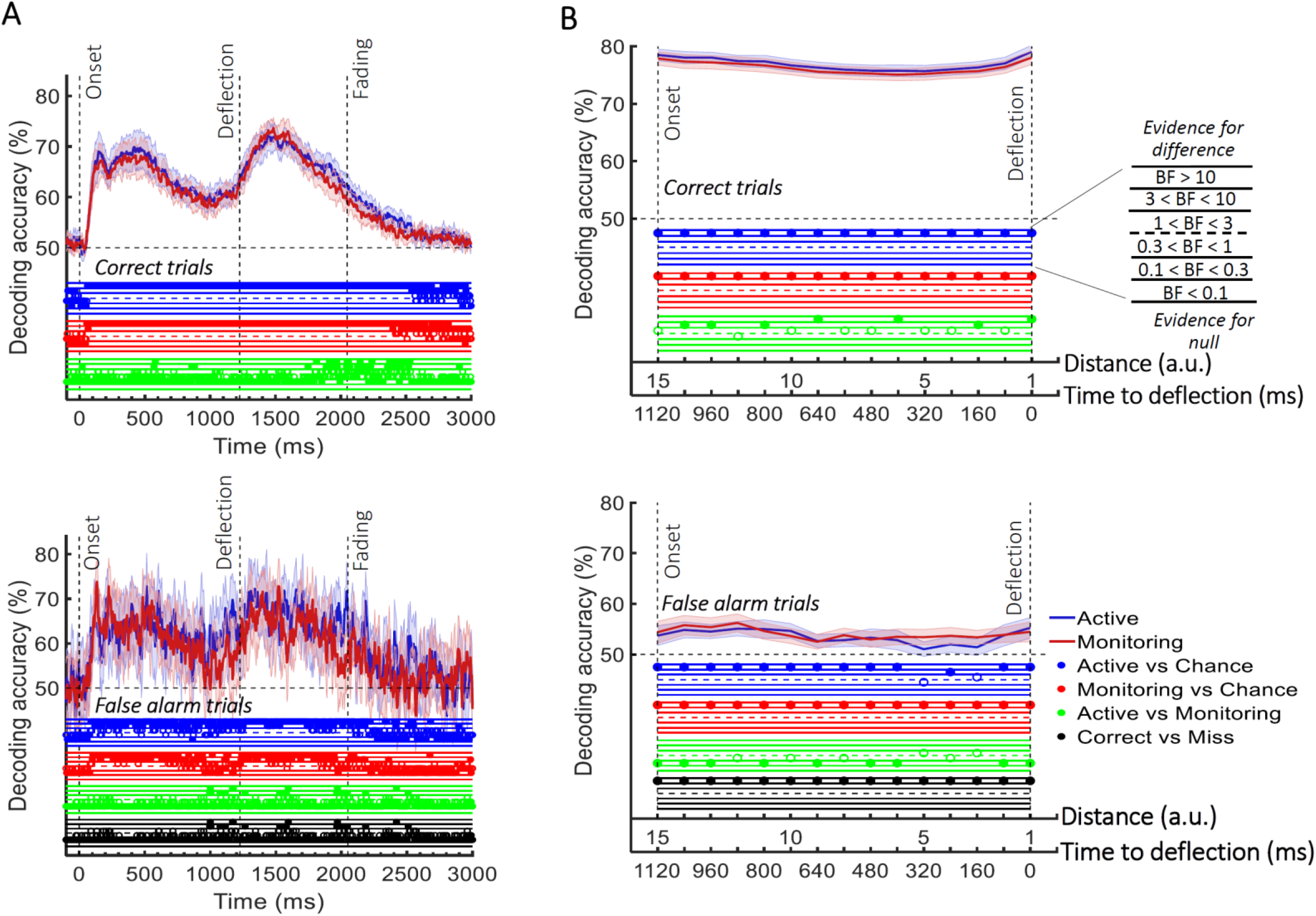
Decoding of information on *correct* vs. *false alarm* trials. (A) Decoding of *direction of approach* information (less task-relevant). (B) Decoding of *distance to object* information (most task-relevant). The horizontal dashed lines refer to chance-level decoding. Top row: Decoding using correct trials; Bottom row: Decoding using false alarm trials. In both top and bottom rows, the classifiers were trained on correct trials and tested on *correct* and *false alarm* trials, respectively. Thick lines show the average across participants (shading 95% confidence intervals). Vertical dashed lines indicate critical events in the trial. Bayes Factors are shown in the bottom section of each graph: Filled circles show moderate/strong evidence for either hypothesis and empty circles indicate insufficient evidence. They show the results of Bayes factor analysis when evaluating the difference of the decoding values from chance for Active (blue) and Monitoring (red) conditions separately, the comparison of the two conditions (green) and the comparison of correct and miss trials (black). Note that for the comparison of correct and miss trials, Active and Monitoring conditions were averaged separately.

### Supplementary Figure 3 shows the analysis of eyetracking data using the same decoding methods as for the neural data

**Supplementary Figure 3.**
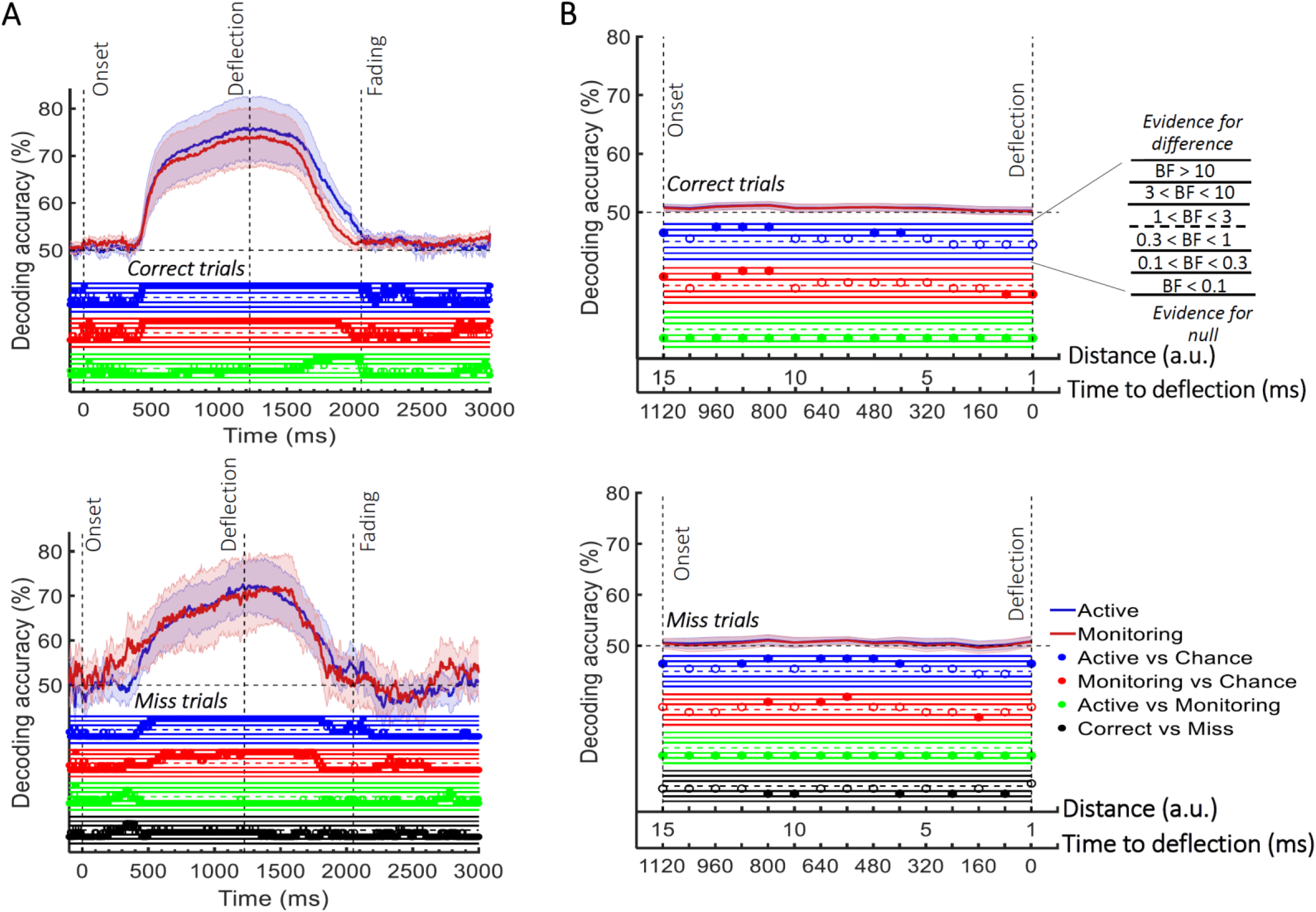
Decoding of information about the dot motion using the eye-tracking data. (A) Decoding of *direction of approach* information (less task-relevant). (B) Decoding of *distance to object* information (most task-relevant). The horizontal dashed lines refer to chance-level decoding. Top panels: Decoding using correct trials; Bottom panels: Decoding using miss trials. In both top and bottom panels, the classifiers were trained on *correct* trials and tested on (left out) *correct* and all *miss* trials, respectively. Thick lines show the average across participants (shading 95% confidence intervals). Vertical dashed lines indicate critical events in the trial. Bayes Factors are shown in the bottom section of each graph: Filled circles show moderate/strong evidence for either hypothesis and empty circles indicate insufficient evidence. They show the results of Bayes factor analysis when evaluating the difference of the decoding values from chance for Active (blue) and Monitoring (red) conditions separately, the comparison of the two conditions (green) and the comparison of correct and miss trials (black). Note that for the comparison of correct and miss trials, Active and Monitoring conditions were averaged separately.

